# Distinct roles and requirements for *Ras* pathway signaling in visceral versus somatic muscle founder specification

**DOI:** 10.1101/347526

**Authors:** Yiyun Zhou, Sarah E. Popadowski, Emily Deustchman, Marc S. Halfon

## Abstract

Pleiotropic signaling pathways must somehow engender specific cellular responses. In the *Drosophila* mesoderm, *Ras* pathway signaling specifies muscle founder cells from among the broader population of myoblasts. For somatic muscles, this is an inductive process mediated by the ETS-domain downstream Ras effectors Pointed and Aop (Yan). We demonstrate here that for the circular visceral muscles, despite superficial similarities, a significantly different specification mechanism is at work. Not only is visceral founder cell specification not dependent on Pointed or Aop, but *Ras* pathway signaling in its entirety can be bypassed. Our results show that de-repression, not activation, is the predominant role of *Ras* signaling in the visceral mesoderm and that accordingly, *Ras* signaling is not required in the absence of repression. The key repressor acts downstream of the transcription factor Lameduck and is likely a member of the ETS transcription factor family. Our findings fit with a growing body of data that point to a complex interplay between the *Ras* pathway, ETS transcription factors, and enhancer binding as a critical mechanism for determining unique responses to *Ras* signaling.

**SUMMARY:** A fundamentally different mechanism is shown for how *Ras* signaling governs cell fate specification in the *Drosophila* somatic versus visceral mesoderms, providing insight into how signaling specificity is achieved.

## INTRODUCTION

Embryonic development requires that individual cells within a field acquire new, distinct fates. The *Drosophila* larval musculature has emerged as an exemplary system for investigating this process, revealing important insights into the acquisition of developmental competence, progressive determination of cell fate, and the integration of multiple signals at the transcriptional level. It has proven to be a particularly effective model for understanding how specific outcomes can be obtained from the activation of the receptor tyrosine kinase (RTK)/Ras/mitogen activated protein kinase (MAPK) signaling cascade (Halfon et al., 2000), a highly pleiotropic pathway involved in numerous developmental processes and dysregulated in a wide set of developmental disorders and cancers (Imperial et al., 2017, Schlessinger, 2000, Tidyman and Rauen, 2009).

In the somatic (body wall) musculature, which has been studied the most extensively, individual syncytial muscle fibers develop via the fusion of two cell types drawn from an initially undifferentiated pool of myoblasts within the stage 10-11 (mid-embryogenesis) mesoderm: a single “founder cell” (FC; itself the product of the asymmetric division of a muscle “progenitor”) and one or more “fusion competent myoblasts” (FCMs; Fig. 1) (for review see Dobi et al., 2015). Whereas FCMs are uncommitted, FCs are induced with specific identities. FCMs fuse with FCs, with the resulting syncytium maintaining the identity provided by the FC. FC specification is thus the critical step in muscle development as the FC genetic program controls muscle size, orientation, expression of cell-surface proteins, and the like.

**Figure 1:**
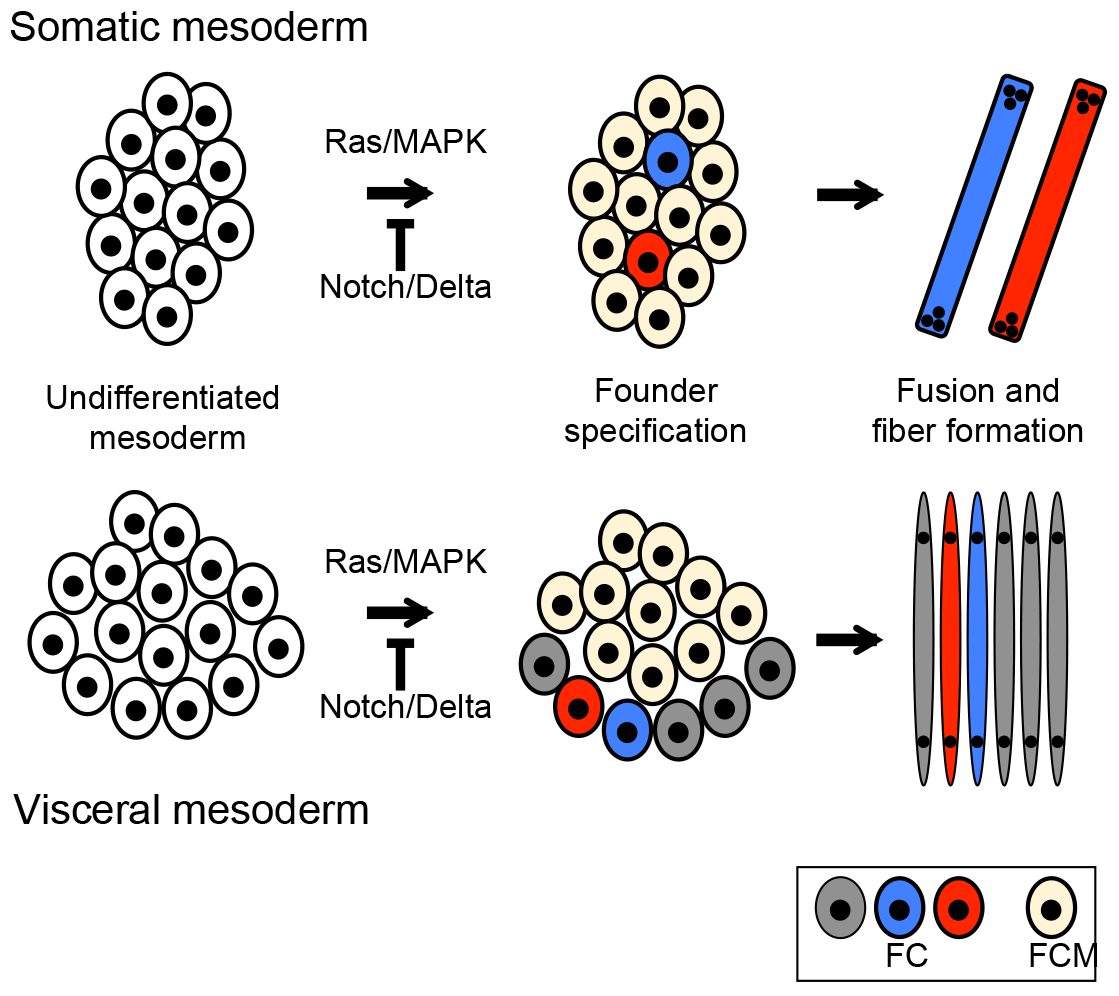
Overview of *Drosophila* muscle development. In both the somatic mesoderm (top) and trunk visceral mesoderm (bottom), initially equivalent myoblasts (left panel) are fated to become either muscle founder cells (FCs; middle panel, gray, red, and blue) or fusion competent myoblasts (FCMs; middle panel, yellow). FCs have specific identities, represented here by different colors, conferred by the activity of “identity genes” active in the FCs. FCMs fuse with FCs to generate individual muscle fibers (right panel), with each fiber maintaining the fate provided by the founder cell.

While multiple signaling pathways, including the Wg (Wnt) and Dpp (BMP) pathways, are integrated to specify particular muscle fates, the key event in muscle determination is MAPK activation via RTK/Ras pathway signaling (Buff et al., 1998, Carmena et al., 2002, Carmena et al., 1998, Halfon et al., 2000). In the somatic mesoderm the relevant receptors are the *Drosophila* EGF and FGF receptor homologs. Cells which are competent to respond to Ras/MAPK signaling are induced as an equivalence group and subsequently restricted by lateral inhibition (mediated by Notch-Delta signaling) to a single muscle progenitor.

These events have been studied in detail at the molecular level in the context of the muscle identity gene *even skipped (eve).* A 300 bp transcriptional enhancer directly integrates the inductive Ras/MAPK signaling with a combination of additional signal-derived and tissue-specific TFs to activate *eve* expression (Halfon et al., 2000). The Ras/MAPK effector is the ETS-domain TF Pnt, which binds the enhancer at up to eight distinct sites (Boisclair Lachance et al., 2018, Halfon et al., 2000). In the absence of induction these sites are bound by the ETS-domain repressor Aop (also known as Yan) (Halfon et al., 2000, Webber et al., 2013, Boisclair Lachance et al., 2018). Recent evidence suggests that Pnt bound at these or other sites may also contribute to repression in the absence of MAPK activation (Webber et al., 2018). Importantly, experiments have shown that induction trumps repression: in the absence of both Pnt and Aop binding, there is no gene activation (Halfon et al., 2000 and unpublished data). Ectopic activation of the Ras/MAPK pathway leads to ectopic FC formation in all competent cells, at the expense of FCMs; this has been demonstrated at the level of the receptor tyrosine kinases (activated EGFR and FGFR), of Ras (activated Ras), and of the effector (activated Pnt) (Carmena et al., 1998, Halfon et al., 2000).

We focus here on the circular visceral muscle fibers, which surround the midgut and develop from the trunk visceral mesoderm (for simplicity, we will refer to these simply as “visceral muscle” and “visceral mesoderm,” respectively). These muscle fibers appear to develop similarly to the somatic muscles (Fig. 1), with the exception that they are only binucleate and it is unclear whether there is a “muscle progenitor” cell specified prior to visceral FC specification (Martin et al., 2001). As with somatic FCs, visceral FC specification occurs following MAPK activation—here via the single signaling pair of the Anaplastic lymphoma kinase (Alk) receptor tyrosine kinase and its ligand Jellybelly (Jeb)—and just as for the somatic musculature, ectopic activation of the Ras/MAPK pathway causes presumptive FCMs to be re-specified as FCs (Fig. 2B) (Englund et al., 2003, Lee et al., 2003, Weiss et al., 2001). However, the details of these events have not been established.

**Figure 2:**
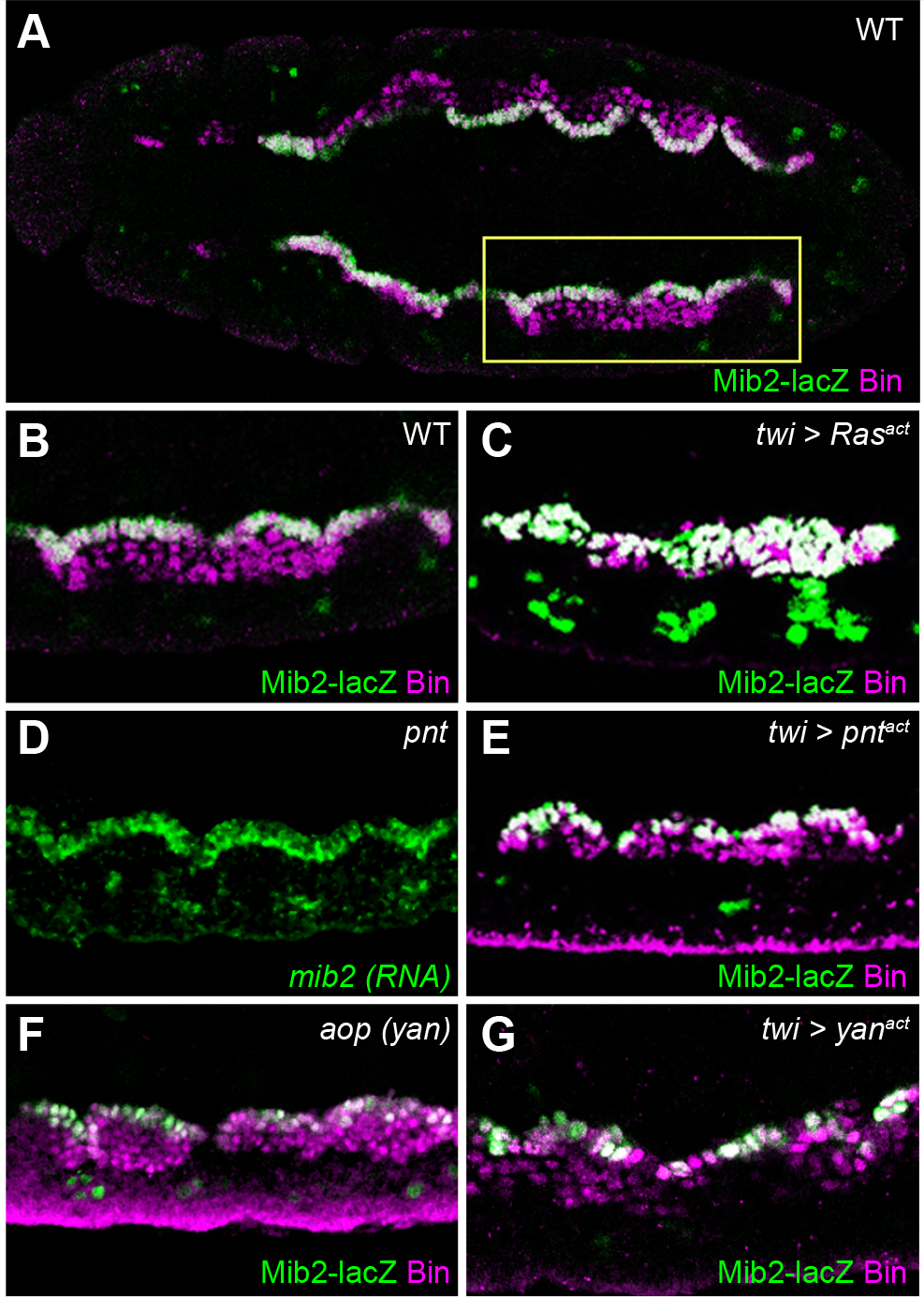
*mib2* expression responds to *Ras* signaling but not to *pnt* or *aop*. All panels show stage 11 embryos stained for expression of Mib2 (using the *mib2_FCenhancer* LacZ reporter, green) and the pan-visceral-mesoderm marker Biniou (Bin, magenta). The exception is panel C, which shows only *mib2* RNA by means of in situ hybridization. (A) Wild type embryo depicted ventral side up and anterior to the left. The yellow box marks the representative segments shown in panels B-G. (B) Wild-type showing Mib2-lacZ expression confined to a single row of visceral mesodermal cells, the FCs. (C) In Twi-Gal4>UAS-Ras^act^ embryos, Mib2-lacZ expression expands throughout the visceral mesoderm. Bin-negative clusters in the foreground are somatic mesoderm. In contrast, *mib2* and Mib2-lacZ expression remains restricted to a single layer of cells corresponding to the FCs, as in wild type, in both a *pnt* null (D) and an activated *pnt* (E) background. Similarly, Mib2-lacZ expression retains a wild-type pattern in *aop* null (F) and *aop* activated (the constitutively repressing *“yan^act^”;* G) backgrounds.

We now show that despite the apparent similarities between somatic and visceral FC specification, fundamental differences exist with respect to the role of Ras/MAPK signaling in specifying the FC fate. Unlike the positive inductive role for MAPK activity in the somatic mesoderm, in the visceral mesoderm, MAPK activity is instead required to relieve repression of FC fates, and the transcriptional effectors Pnt and Aop do not appear to play a significant role in this process. Moreover, MAPK activity can be dispensed with entirely in the absence of the FCM-specific transcription factor Lameduck (Lmd) or when repressor binding sites are mutated in an FC-specific enhancer for the *mib2* gene. Thus, unlike in the somatic mesoderm, Ras/MAPK signaling is not essential for visceral FC specification. Our results illustrate how similar-appearing developmental processes can result from different underlying molecular mechanisms and provide fresh insights into how unique responses to Ras-pathway signaling are determined within similar cellular and developmental contexts.

## RESULTS

### Visceral founder cell specification is independent of the ETS-domain transcription factors Pnt and Aop

In a previous screen for genes that respond differentially to different Ras-pathway components, we observed that despite responding to RTK and Ras activation, the FC gene *mib2* is not regulated by the Ras effector Pnt in the visceral mesoderm (Leatherbarrow and Halfon, 2009). Expression of both *mib2* RNA and a *mib2-lacZ* reporter gene driven by an FC-specific enhancer *(mib2-FCenhancer)* is normal in *pnt* null mutant embryos (Fig. 2C and Halfon et al., 2011), and expression of a constitutively active form of Pnt (Pnt^act^) has no effect on expression of either the endogenous gene or the reporter (Fig. 2D and Leatherbarrow and Halfon, 2009). Similarly, *mib2* expression in the visceral mesoderm is normal in embryos mutant for the ETS-domain repressor *aop (yan)* (Fig. 2E), and in response to expression of the constitutively-repressing version *“yan^act^”* (Fig. 2F and Halfon et al., 2011). This raised the question of whether this is a *mib2-* specific regulatory effect, or whether these two Ras effectors, which both play a significant role in somatic FC determination, are not required for visceral FC specification.

To test this, we assessed the expression of additional visceral FC and FCM markers in *pnt* null and/or *pnf^act^* backgrounds. Expression of the somatic muscle identity gene *even skipped (eve)* was used as a control (data not shown), as its expression is respectively reduced or expanded in response to *pnt* loss and gain of function. Expression of the FC markers *org-1, kirre* (also known as *dumbfounded (duf*)), and *RhoGAP15B* all appear normal in a *pnt^act^* background, whereas, as reported previously, expression of all three expands with pan-mesodermal expression of activated Ras (Ras^act^) (Fig. 3; Leatherbarrow and Halfon, 2009, Lee et al., 2003). Double-labeling with antibodies to Biniou, a general visceral mesoderm marker (Zaffran et al., 2001), showed that the observed expansion is throughout the visceral mesoderm (Fig. 2B and data not shown). This suggests that non-FCs (i.e., FCMs) have been respecified as FCs, rather than that there has merely been increased FC proliferation. Consistent with this, expression of the FCM marker Lmd behaves in the reciprocal fashion: Lmd expression decreases in the visceral mesoderm with *Ras^act^* expression, but is unaffected in *pnt* null or *pnt^act^* backgrounds (Fig. 3B, E, H and data not shown; Popichenko et al., 2013). Taken together, our results show that while Ras activity is sufficient to induce FC fates throughout the visceral mesoderm, neither *pnt* nor *aop* appear to play a significant role in this process.

**Figure 3:**
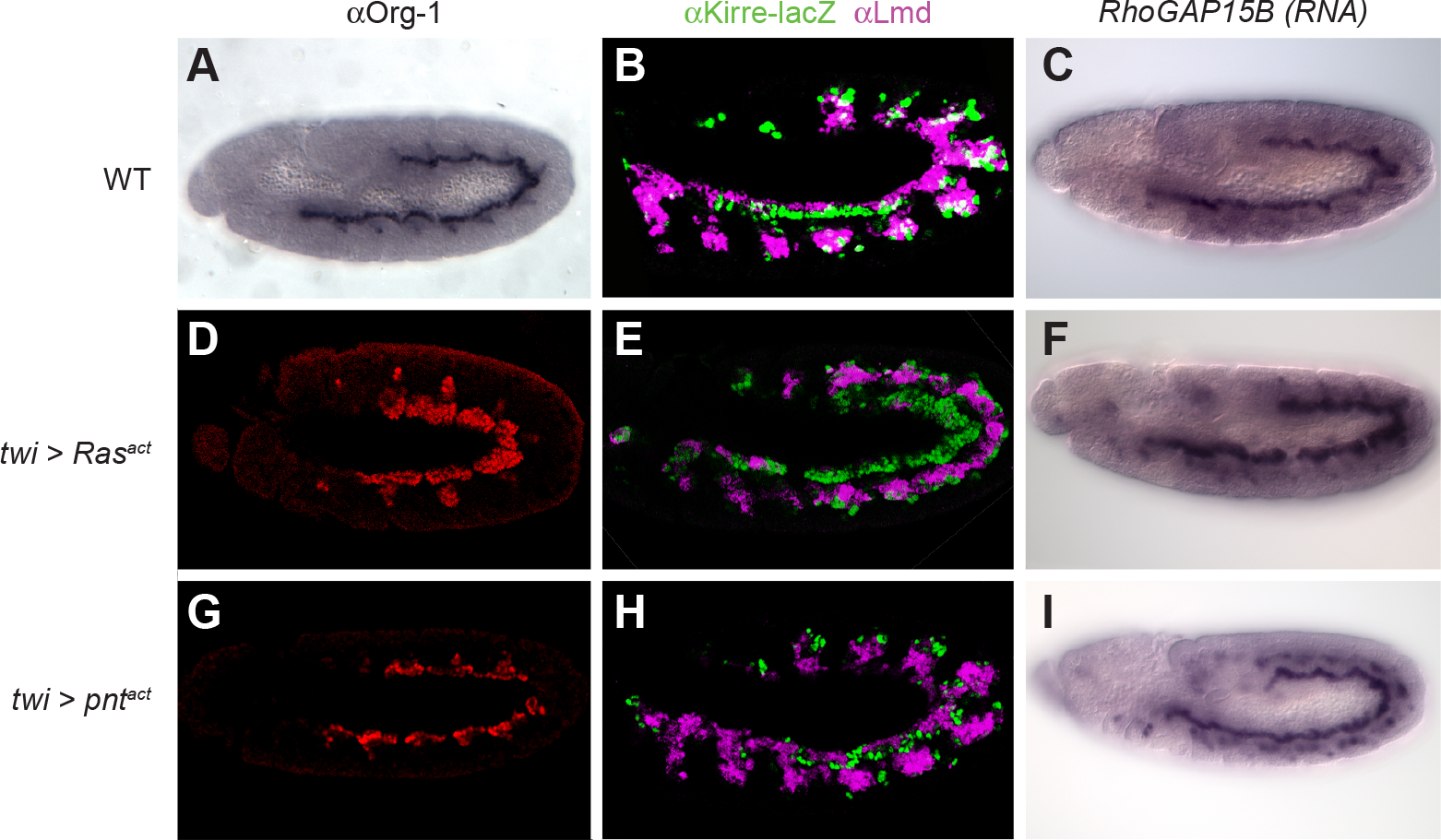
Expression of multiple visceral FC and FCM markers respond to *Ras* but not to *pnt*. Stage 11 embryos that are either wild type (A-C), have pan-mesodermal expression of activated Ras (D-F; *“twi>Ras^act^”*), or have pan-mesodermal expression of activated Pnt (G-I; “*twi>pnt^act^”* were stained for FC and/or FCM markers. Consistent with results assessing *mib2* expression, Ras activation led to increased FC and decreased FCM populations, while Pnt activation had no effect. Panels A, D, and G show expression of Org-1, an FC marker; B, E, and H depict the FC marker *kirre^rp298^~^PZ^*, an enhancer-trap in the *kirre (duf)* locus (green), and FCM marker Lmd (magenta); and C, F, and I picture in situ hybridization to RhoGAP15B RNA. All embryos are oriented dorsal up and anterior to the left.

### Visceral founder cell expression of *mib2* is repressed through ETS-type binding sites in the *mib2* FC enhancer

Although *mib2* expression in the visceral mesoderm is not dependent on either *pnt* or *aop*, the *mib2_FCenhancer* enhancer was identified in part based on the presence of ETS-type, putative Pnt binding sites (Philippakis et al., 2006). We showed previously that mutation of a set of seven ETS sequences in this enhancer caused an expansion of reporter gene expression driven by the mutated enhancer (Fig. 4E, F and Halfon et al., 2011). Like the expression observed with activation of the *Ras* pathway, the expanded reporter gene expression extends throughout the visceral myoblast population as marked by Bin expression (data not shown). Interestingly, reporter gene expression is stronger in the FCs than in the rest of the myoblasts (Fig. 4G, Fig. S1A and data not shown). Activated Ras expression restores full-strength reporter activity similar to what is observed with the wild-type enhancer (Fig. 4H, Fig. S1B and data not shown). However, as expected, the same disparity in reporter expression between FCs and non-FCs as seen in the wild-type background is seen with activated Pnt, which by itself does not lead to expanded *mib2* expression (Fig. 4I, Fig. S1C and data not shown).

**Figure 4:**
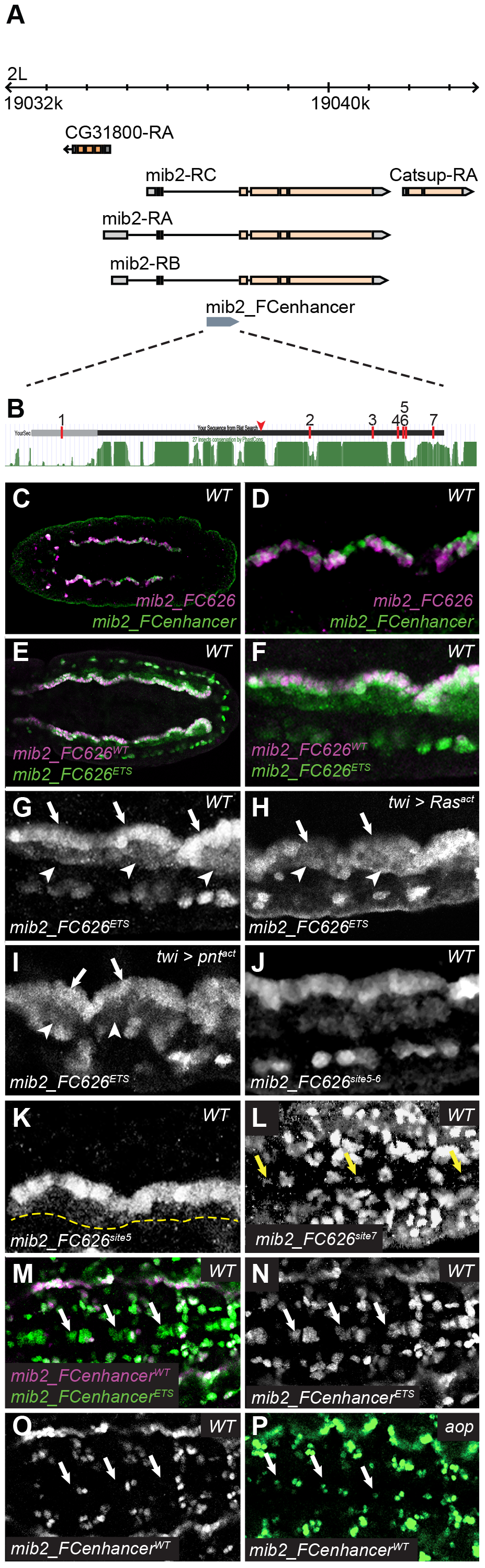
Mutagenesis of the *mib2* FC enhancer reveals repression acting via ETS binding sites. (A) Schematic of the *mib2* locus. The location of the intronic *mib2_FCenhancer* regulatory sequence is indicated in gray. (B) The *mib2_FCenhancer* regulatory sequence, with conservation shown below in green. The gray portion of the sequence is deleted in the *mib2_FC626* constructs. A red arrowhead marks the location of the region deleted in the inactive 413 bp 5’ deletion. Red bars numbered 1-7 indicate the positions of the tested ETS binding sites. Conservation track shows the 27-insect PhastCons conservation from the UCSC Genome Browser. (C) The shorter *mib2-FC626* enhancer (magenta) has activity indistinguishable from the original *mib2_FCenhancer* enhancer (green); a higher magnification view can be seen in (D). (E, F) Mutation of the 6 ETS binding sites in the *mib2-FC626* enhancer (“*mib2-FC626*^ETS^, green) causes an expansion of reporter gene expression throughout the visceral mesoderm. Expression in the FCs is stronger than the ectopic FCM-domain expression (G). In contrast, pan-mesodermal Ras activation causes similar ectopic expression, but reporter gene levels are consistent throughout the visceral mesoderm (H). Expression of activated Pnt, however, resembles the expression seen in a wild-type background (I). (J) Mutation of ETS sites 5 and 6 *(“mib2-FC626^site5^~^6^”)* causes reporter gene expression to expand into the FCM domain, but the expanded expression is considerably weaker than that seen with the full 6-site mutation (compare with panel G). (K) Mutation of site 5 alone *(“mib2-FC626^site5^”)* also causes a weak reporter gene expansion. The yellow dotted line indicates the border of the FCM domain, as assessed by two-color imaging for the pan-visceral mesoderm marker Biniou (not shown). (L) Mutation of site 7 (“*mib2-FC626^sit7^”)* has no effect in the visceral mesoderm (not shown), but leads to ectopic reporter gene expression in the midline of the ventral nerve cord (arrows). Similar ectopic expression is observed when all ETS sites are mutated (M, N, arrows; compare to the same locations marked by arrows with the wild-type enhancer in panel O). (P) Similar ectopic reporter gene expression in the ventral midline is also seen with the wild-type enhancer in a *aop* mutant background (arrows).

The expanded reporter gene expression observed upon mutation of the ETS sequences suggested that rather than being required for positively activating *mib2* expression—as expected based on analogy to the requirement for *Ras* pathway signaling mediated by ETS-family transcription factors in the somatic mesoderm (Halfon et al., 2000)—*mib2* is repressed via transcription factor binding at these sites. In order to better understand the nature of this repression, we decided to characterize the *mib2* regulatory sequences more thoroughly.

Using sequence conservation with other *Drosophila* species as a guide, we first tested reporter gene activity with a truncated version of the *mib2_FCenhancer* containing a 5’ 120 bp deletion (Fig. 4B, Fig. S2). The deleted region includes one of the putative Pnt binding sites previously mutated (“site 1”, Fig. 4B, Fig. S2), as well as a non-canonical Pnt site suggested by protein binding microarray experiments (Fig. S2, “siteN”; personal communication from Alan Michelson). The resulting “*mib2_FC626*’ enhancer has activity identical to the longer *mib2_FCenhancer* (Fig. 4C, D), responds to *Ras^act^* and *Pnt^act^* ectopic expression in the same manner (Fig. S1E, F and Leatherbarrow and Halfon, 2009), and shows a similar Ras^act^-like expansion of reporter gene expression when the remaining six ETS-type sequences are mutated (construct *“mib2_FC626^ETS^*; Fig. 4E, F, G). In contrast, a more extensive 5’ 413 bp deletion (Fig. 4B arrowhead, Fig. S2, gray arrow) leads to a complete loss of visceral mesoderm activity and only a limited residual expression elsewhere (data not shown). As the *mib2_FC626* enhancer behaves in all aspects like the original *mib2_FCenhancer*, we used this shorter sequence as a template for further characterization of *mib2* regulation.

We mutated the six remaining ETS-type sequences individually to determine which putative binding sites were responsible for the expanded reporter gene expression (as sites 5 and 6 are close together, we treated them initially as a single site5-6). Expanded reporter gene expression was observed only with the site5-6 paired mutation (Fig. 4J, Fig. S3). We therefore further dissected this pair to test its component individual sites. Mutation of site 5 led to expansion of *mib2_FC626* enhancer activity throughout the visceral mesoderm (Fig. 4K), while site 6 by itself had a barely observable phenotype with expanded expression almost indistinguishable from background (Fig. S1G). The expanded expression observed with the single site 5 mutation was notably weaker than that observed with the paired site5-6 mutation, which itself had weaker expression than the *mib2_FC626^ETS^* six-site-mutated enhancer (Compare Fig. 4G, J, and K). While site 5 therefore appears to be the most critical site contributing to expanded *mib2_FC626* expression, the stronger expression seen when multiple sites are mutated suggests that these other sites are functional as well and contribute to overall enhancer activity. Consistent with their more essential roles, site 5 is the most highly conserved of the six ETS sites in the *mib2_FC626* sequence, followed by site 6 (Fig. S2B).

In addition to expanded visceral mesoderm expression, the *mib2_FC626^ETS^* construct drives ectopic reporter gene expression in the midline of the ventral nerve cord (Fig. 4M-O and Halfon et al., 2011). This ectopic midline expression is also observed with the *mib2_FC626^site7^* construct (Fig. 4L), but not with any of the other single-site enhancer mutations. Similar ectopic *mib2* expression is observed in *aop* null mutant embryos (Fig. 4P), suggesting that while Aop does not regulate *mib2* in the visceral mesoderm, it may act via this site to repress *mib2* in the nervous system.

### The de-repressed *mib2* enhancer does not require *Ras* pathway signaling for its activity

In the somatic mesoderm, we showed previously that *Ras* pathway activity is absolutely required for FC gene expression; for example, in the absence of both *pnt* and *aop* expression the *eve_MHE* FC enhancer is inactive, and mutation of the common ETS-type Pnt and Aop binding sites eliminates enhancer activity (Halfon et al., 2000). However, the de-repression of the *mib2* visceral FC enhancer seen with ETS site mutation suggested that *Ras* activity, normally not present in the FCM population into which *mib2* reporter gene expression expands, might be dispensable when the *mib2* enhancer is de-repressed. To test this, we analyzed the activity of the wild-type and mutated *mib2_FC626* enhancers in a *jeb* null background, which lacks *Ras* signaling in the visceral mesoderm. Whereas the wild type *mib2_FC626* enhancer is inactive in the visceral mesoderm of *jeb* null embryos (Fig. 5A), the *mib2_FC626^ETS^* mutated enhancer is expressed throughout the visceral mesoderm just as in a wild type background (Fig. 5B, arrows). Staining for the activated, di-phosphorylated form of MAPK confirmed that no signaling was present in the *jeb* null background (Fig. 5D). Unlike in the somatic mesoderm, therefore, *Ras* signaling is not required for expression of a visceral FC gene in the absence of ETS-site mediated repression.

**Figure 5:**
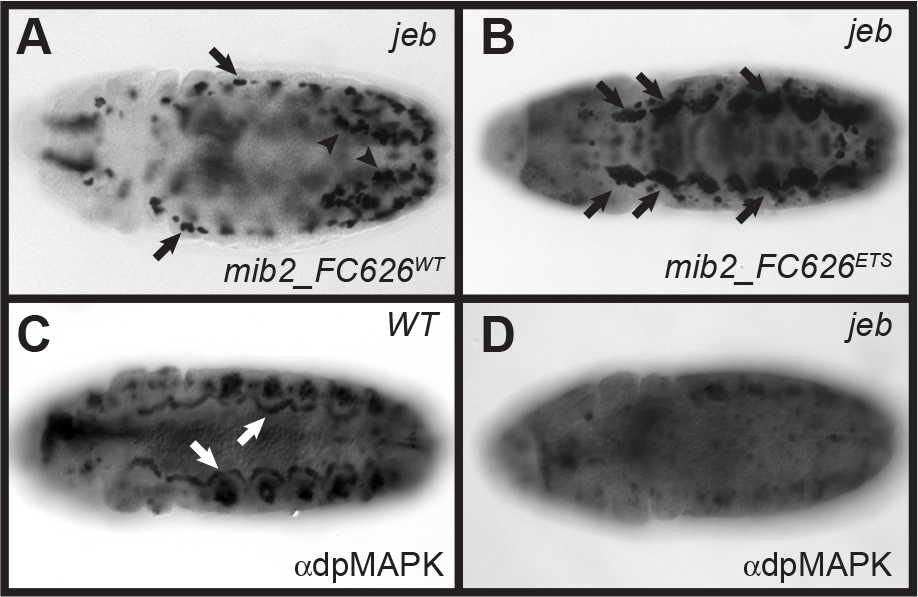
The mutated *mib2* enhancer is active even in the absence of *Ras* signaling. (A) The wild-type (“*mib2-FC626^WT^”)* and (B) ETS-site mutated (“*mib2-FC626^ETS^”) mib2* enhancers were crossed into a *jeb* null background that lacks *Ras* signaling in the visceral mesoderm and assessed for reporter gene expression. Trunk visceral mesoderm expression is absent for the wild-type enhancer, with only somatic mesoderm (arrows) and caudal visceral mesoderm (arrowheads) activity still present. However, robust visceral mesoderm activity resembling that seen in a wild-type background is observed with the mutated enhancer (B, arrows). This expression resembles that seen with the wild-type enhancer following ectopic Ras activation (compare with Figure 2B). (C) Staining for activated MAPK (dpMAPK) shows crescents of visceral mesoderm expression in wild type embryos (arrows), which are absent in *jeb* null embryos (D). Embryos are oriented ventral up and anterior to the left.

### Visceral FC specification can occur in the absence of *Ras* pathway signaling

The expanded expression of an FC gene throughout the visceral mesoderm we observed in the case of the de-repressed *mib2_FC626* enhancer is reminiscent of what has been observed upon loss of function of the FCM transcriptional activator Lmd. Popichenko et al. (2013) showed that in *lmd* null embryos, FC markers such as *org-1, hand*, and *kirre* expand throughout the visceral mesoderm, with reciprocal loss of FCM genes such as *Vrp1.* This resembles the phenotype observed with Ras activation. In a similar fashion, *lmd* mutation leads to the conversion of a small subset of FCMs to adult muscle precursor and pericardial cells (Sellin et al., 2009). However, these phenotypes are in sharp contrast to what is observed for the bulk of the somatic mesoderm, where *lmd* loss of function has no effect on FC specification (Duan et al., 2001, Ruiz-Gomez et al., 2002). Interestingly, the FCM-specific gene *sticks-and-stones (sns)* remains expressed in the *lmd* visceral mesoderm, suggesting that the observed FCMx→FC conversion is incomplete (Estrada et al., 2006, Ruiz-Gomez et al., 2002). Given our results with the *mib2* enhancer, we wondered if *lmd* loss-of-function induced FCM→FC respecification could also occur in the absence of Ras pathway activity. Therefore, we tested *jeb;lmd* double mutant embryos for a range of FC markers including *org-1, mib2*, and *RhoGAP15B*, which are expressed in all FCs, as well as *connectin (con)* and *wingless (wg)*, which are expressed in only a subset of FCs (Bilder and Scott, 1998). In all cases, *lmd* was fully epistatic to jeb: whereas in *jeb* null embryos no FC markers are expressed, expression in *jeb;lmd* embryos consistently resembles *lmd* alone, in most cases with expanded FC expression (Fig. 6). Surprisingly, *mib2* and *RhoGAP15B*, which expand throughout the visceral mesoderm both with Ras activation and upon mutation of the *mib2* FC enhancer, do not appear to have expanded expression in the *lmd* background when assayed by in situ hybridization (Fig. 6B, 6F). This may be indicative of an incomplete conversion of FCMs to FCs, consistent with the maintenance of *sns* expression in the FCM region reported previously. Likewise, Wg expression also only expands to a few additional cells and not throughout the entire width of the anterior PS8 visceral mesoderm (Fig. 6S). On the other hand, we found that the *mib2_FC626* reporter construct does show fully expanded expression in *lmd* embryos (Fig. 6B’). As expression driven by the *mib2_FC626* enhancer appears identical to endogenous *mib2* expression in all other contexts we have examined, it may be that the lack of expanded *mib2* expression observed in *lmd* embryos simply represents a failure of detection, given that, similar to what we saw with the reporter construct in other backgrounds, reporter gene expression in the FCM region is weaker than that in the native FC region (Fig. S1D). Consistent with this interpretation, we see a modest widening of *mib2* expression in the *jeb;lmd* embryos (Fig. 6D). Importantly, regardless of the exact degree of expanded expression due to *lmd* loss of function, FC expression of all tested genes is clearly rescued in the *jeb;lmd* background, demonstrating the ability for *Ras* signaling to be bypassed in the absence of *lmd* expression.

**Figure 6:**
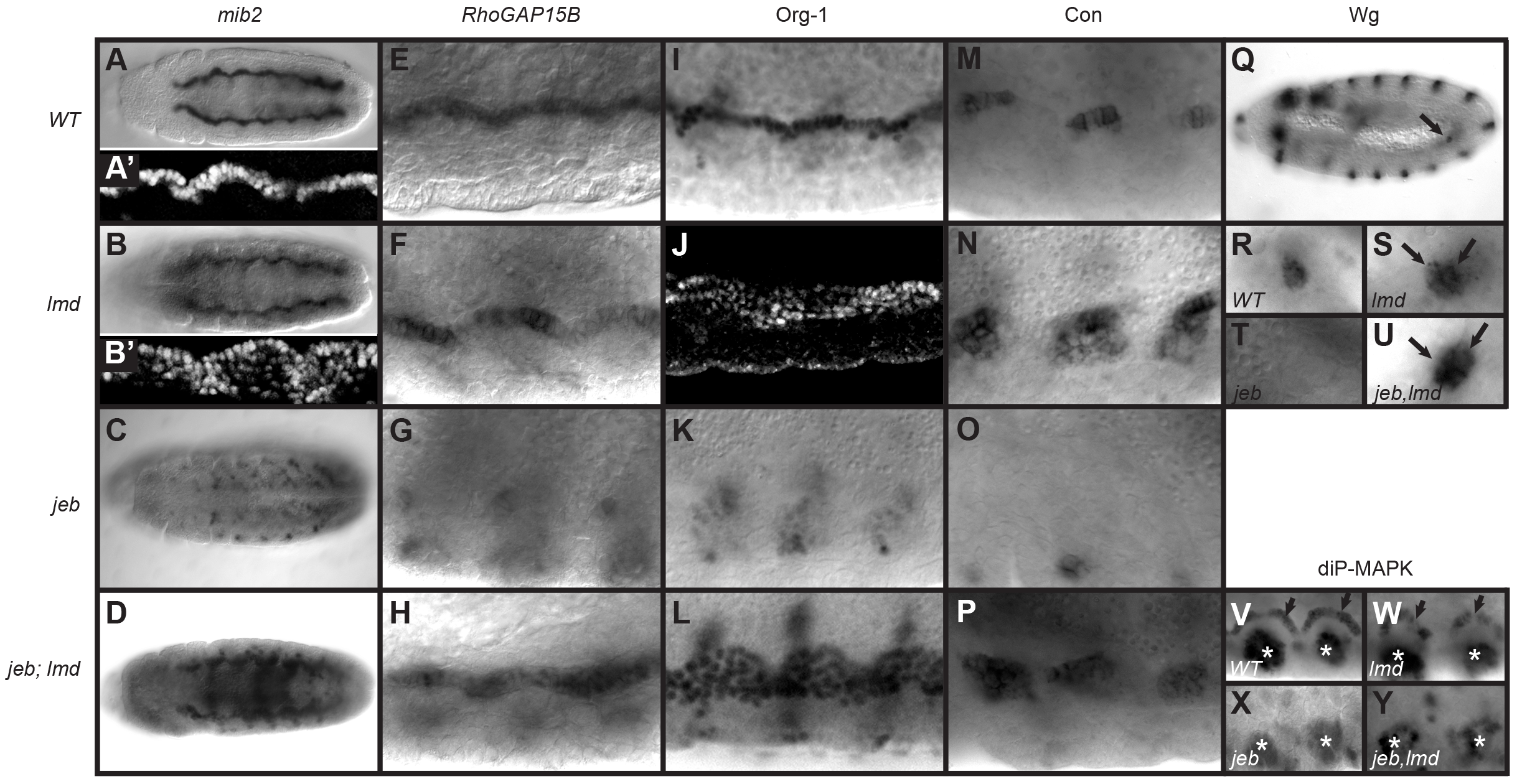
FC gene expression is expanded in *lmd* and *jeb,-lmd* mutant embryos. Expression of the FC genes *mib2* (A-D), *RhoGAP15B* (E-H), *Org-1* (I-L), *con* (M-P), *wg* (Q-U), and activated *MAPK* (V-Y) was assessed in wild type, *lmd, jeb*, and *jeb;lmd* double mutant embryos. Embryos in panels A-P and V-Y are oriented ventral up and anterior to the left and are shown as either whole embryos or as three-segment closeups (two segments in V-Y). Panel Q is oriented with ventral to the bottom and anterior to the left; the arrow points to the region shown in close up in panels R-U. (A) In situ hybridization for *mib2* RNA. (A’) Reporter gene expression driven by the *mib2_FC626* enhancer. (B) In the *lmd* mutant background, *mib2* RNA expression resembles wild type, but the reporter construct (B’) has expanded expression similar to that seen with Ras activation or ETS site mutation. (C) Visceral mesoderm expression of *mib2* is absent in *jeb* embryos but (D) is restored and mildly expanded in the *jeb;lmd* background. *RhoGAP15B* expression (E) does not show expansion in *lmd* embryos (F), but is likewise absent in *jeb* (G) and restored in a *jeb;lmd* (H) mutant background. (I-L) Org-1 expression expands in *lmd* (J), is lost in *jeb* (K), and is restored in *jeb;lmd* (L) embryos. (M-P) The same is true for Con, although expression remains confined to its wild-type anterior-posterior domain. (Q) Wg is expressed in a single visceral muscle FC in the wild type stage 11 embryo (arrow). Higher magnification views reveal that the cell number is increased in *lmd* (S, arrows) and *jeb;lmd* (U, arrows) mutant embryos, but Wg expression is absent in the *jeb* null background (T). (V-Y) Staining for activated (di-P) MAPK confirms that MAPK activation is normal in *lmd* embryos (W) but absent in the *jeb* (X) and *jeb;lmd* (Y) visceral mesoderms. Arrows indicate visceral mesoderm expression, asterisks mark MAPK activation in the tracheal pits.

To ensure that the rescue of FC specification observed in *jeb;lmd* mutants is not the result of a cryptic *Ras* signaling pathway activated by loss of *lmd*, we checked for the presence of activated MAPK in the double-mutant embryos. As expected when *jeb* is absent, no activated MAPK is observed, regardless of presence or absence of *lmd* (Fig. 6X, Y).

## DISCUSSION

Both somatic and visceral muscle development require as an initial step the specification of individual muscle founder cells from within the general myoblast pool. Superficially, the process appears alike for both tissues: FCs fail to form in the absence of RTK/Ras/MAPK signaling, and ectopic activation of the *Ras* pathway causes FCMs to be respecified as FCs. A striking aspect of our current results is that these seemingly similar events are brought about in a mechanistically opposite fashion in the somatic versus visceral mesoderms. Our work therefore serves to underscore how common developmental outcomes can derive from dramatically different gene regulatory mechanisms.

In the somatic mesoderm, it has been well-established that positive induction via Ras/MAPK signaling is essential for specifying FC fates (Buff et al., 1998, Carmena et al., 2002, Carmena et al., 1998, Halfon et al., 2000). In the visceral mesoderm, however, repression has primacy over induction. We demonstrate here that Ras/MAPK signaling acts in presumptive FCs to relieve repression of the FC fate, while Popichenko et al. (2013) previously established that it serves to prevent activation of FCM genes. The primary activator of FCM genes is Lmd, which prior to FC specification is expressed in all visceral myoblasts (Ruiz-Gomez et al., 2002, Popichenko et al., 2013). Ras/MAPK signaling in FCs causes phosphorylation of Lmd followed by its export from the nucleus and its degradation, preventing it from activating FCM-specific genes such as *Vrp1* (Popichenko et al., 2013). What happens at FC gene loci, however, had not previously been determined. Our results with the *mib2* enhancer demonstrate that FC genes can be activated in the absence of Ras/MAPK signaling, through loss of repressor binding at the enhancer.

### A model for FC fate specification

The simplest model, taking into account our results and those of Popichenko et al. (2013), would be for Lmd to function as both the FC gene repressor and the FCM gene activator; loss of Lmd binding following Ras/MAPK signaling would thus simultaneously de-repress FC genes while halting activation of FCM genes. Although Lmd is typically viewed as an activator, some evidence suggests that it may also be capable of transcriptional repression (Cunha et al., 2010). However, chromatin immunoprecipitation studies have repeatedly failed to detect appreciable Lmd binding in the *mib2* enhancer region (Busser et al., 2012, Cunha et al., 2010), and the *mib2* enhancer lacks good candidate Lmd binding sites—particularly in the critical site5-site6 region (MSH, unpublished results)—even when surveyed using a range of binding motifs derived from multiple sources (Busser et al., 2012, Nitta et al., 2015, Zhu et al., 2011).

We therefore favor a basic model in which Lmd serves an activator of the FC gene repressor, such that loss of Lmd leads to loss of repressor activity in FCs and subsequent expression of FC genes (Fig. 7). In wild type embryos, the main role of MAPK signaling is thus to cause phosphorylation and degradation of Lmd, whereas in *lmd* mutant embryos, MAPK signaling becomes irrelevant as Lmd is already absent. The repressor could also be a direct target of MAPK, leading to its rapid displacement upon the onset of MAPK signaling (e.g., similar to what happens with Aop at other loci (Rebay and Rubin, 1995)). This would be consistent with the rapid timecourse of FC fate specification following MAPK activation. We surmise that there are also additional, still unknown FC gene activators whose activity may or may not be MAPK dependent. Tests of these various refinements to the basic model will require identification of the FC gene repressor.

**Figure 7:**
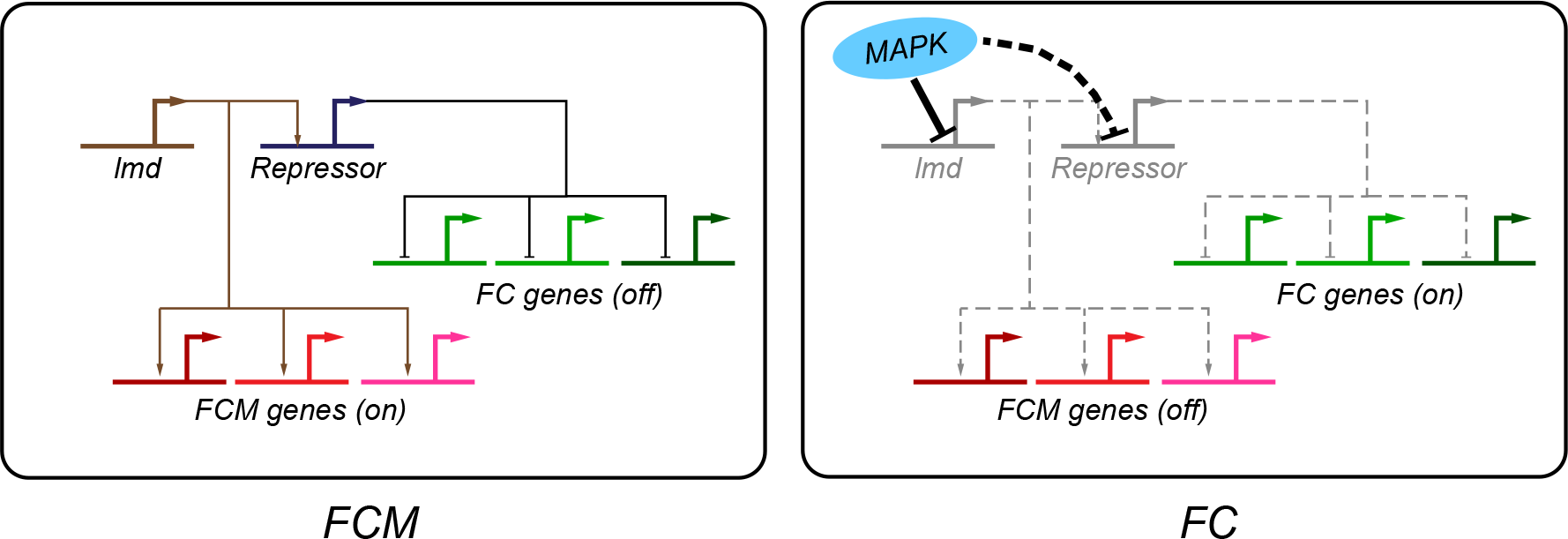
A model for visceral founder cell specification. In FCMs (left), Lmd activates FCM-specific genes as well as an FC-gene repressor, which keeps FC-specific genes shut off. In FCs (right), activation of MAPK leads to the degradation of Lmd, preventing activation of both the FCM genes and the FC-gene repressor. MAPK may also act directly on the FC-gene repressor (dotted line). Loss of repression allows for expression of the FC genes, possibly in conjunction with additional activators (not pictured).

In the somatic mesoderm, the repressor Tramtrack69 (Ttk69) appears to play a role as an *lmd*-dependent FC gene repressor similar to what we posit here for visceral FC fate repression. *ttk69* expression is activated downstream of *lmd* in FCMs, where it represses the transcription of FC genes (Ciglar et al., 2014). In FCs, Ttk69 is likely post-translationally degraded in a Ras-dependent manner (Ciglar et al., 2014, Li et al., 1997), relieving repression of FC genes concurrent with Ras-dependent relief of Aop-mediated repression and induction via Pnt and/or other activators. However, different mechanisms appear to be at work in the visceral mesoderm. Although *ttk69* mutants do display some altered visceral mesoderm gene expression (Ciglar et al., 2014), visceral FCs appear to be correctly specified and FC-specific genes such as *mib2* are expressed in the appropriate pattern, without expansion into the FCM field (Ciglar et al., 2014; SEP, unpublished observations).

### A balance of ETS factors?

Given the derepression phenotypes observed on mutation of the ETS sites in the *mib2* enhancer, we favor the likelihood that the relevant repressor is a member of the ETS transcription factor family, either by itself or working redundantly with Pnt and/or Aop. Several other ETS domain proteins exist in *Drosophila* (Chen et al., 1992), although their expression patterns and mutant phenotypes are for the most part not well defined. A role for additional ETS proteins was also previously suggested for the dorsal somatic mesoderm, where *pnt* loss-of-function leads to a partial loss of Eve-expressing FCs, but mutation of ETS binding sites completely eliminates expression driven by the *eve_MHE* enhancer (Halfon et al., 2000). Although our data argue against an absolute requirement for either Pnt or Aop, we cannot rule out a more limited contribution from these factors. Indeed, chromatin IP experiments indicate that Pnt can bind to the *mib2* enhancer region, although it is not known in what cell types (Webber et al., 2018), and *aop* mutants show an effect on *mib2* expression in the ventral midline (Fig. 4P).

Although there is considerable evidence demonstrating the requirement for RTK/Ras/MAPK signaling for somatic FC specification, the molecular details on the mechanisms governing MAPK-dependent activation and repression come mainly from studies of a single transcriptional enhancer, *eve_MHE* (Boisclair Lachance et al., 2018, Halfon et al., 2000, Webber et al., 2018, Webber et al., 2013). It is clear that ETS-factor-dependent activation is essential for the activity of this enhancer, as mutation of the major ETS binding sites renders the enhancer non-functional (Halfon et al., 2000). However, recent studies suggest that instead of an abrupt switch between activation and repression due to mutually exclusive enhancer occupancy by Pnt and Aop, there is a more subtle balance between these transcription factors and their binding to the multiple high-and low-affinity ETS binding sites found in the enhancer (Boisclair Lachance et al., 2018, Webber et al., 2018). Other somatic FC enhancers have not been rigorously tested with respect to ETS-family binding, and it may be that the trade-off between activation and repression differs among them. This would help to explain the results of Buff et al. (1998), who demonstrated that different FCs are specified at different levels of RTK/Ras signaling. One way to achieve such differential sensitivity would be through a mixture of activating versus repressing ETS transcription factors competing for binding at a range of high-and low-affinity sites. Such a mechanism could provide for exquisitely fine-tuned responsiveness to Ras/MAPK signaling, making this an appealing possibility.

In this vein, we note that our detailed molecular insights for visceral FCs again come mainly from the study of a single enhancer, *mib2_FC626.* Here, elimination of the major ETS binding sites leads to increased activity, opposite the situation with the somatic *eve_MHE* enhancer. However, preliminary analysis of other visceral FC enhancers suggests that eliminating ETS binding sites can lead to loss of enhancer function, more similar to what is seen in the somatic musculature (YZ, unpublished observations). Thus while our data from the *mib2_FC626* enhancer as well as from analysis of *lmd* mutants clearly establishes de-repression rather than induction as the major role for *Ras* pathway signaling during visceral FC specification, it may be that the molecular basis for how this signaling is modulated by ETS-family transcription factors at the enhancer level is complex and balanced individually at each FC gene enhancer. Taken together, these plus other recent results (Boisclair Lachance et al., 2018, Webber et al., 2018) point to an elaborate interplay between *Ras* signaling, ETS transcription factors, and subtly tuned binding sites, and highlight the need for detailed molecular studies of a more comprehensive set of both somatic and visceral FC enhancers.

## MATERIALS AND METHODS

### Drosophila strains and genetics

*Oregon-R* was used as the wild type. Mutant stocks are described in FlyBase (Gramates et al., 2017) and include *pnt^Δ88^, aop^1^, lmd^1^*, and *jeb^576^* (Weiss et al., 2001). *mib2_FCenhancer-lacZ* is described in (Philippakis et al., 2006) and the *rp298-lacZ* (FlyBase: *kirre^rp298^~^PZ^*) line was used to assess *kirre (duf)* expression (Nose et al., 1998). Ectopic expression was achieved using the Gal4-UAS system (Brand and Perrimon, 1993) and used lines *Twi-Gal4* (FlyBase: *P{Gal4-* twi.G})(Greig and Akam, 1993), *UAS-Ras1^Act^* (Carmena et al., 1998), *UAS-PntP2VP16* (Halfon et al., 2000), and *UAS-yan^Act^* (Rebay and Rubin, 1995). Mutant lines were rebalanced over *lacZ-* marked balancers to allow for genotyping of embryos.

### Immunohistochemistry and Microscopy

Antibody staining was performed using standard *Drosophila* methods. The following primary antibodies were used: mouse α-β-galactosidase (Promega #Z3783), 1:500; rabbit α-GFP (Abcam ab290), 1:10000; rabbit α-Bin (gift of Eileen Furlong), 1:300; rabbit α-Lmd (gift of Hanh Nguyen), 1:1000; rat α-Org-1 (gift of Manfred Frasch), 1:250; mouse *α*-activated MAPK (diphosphorylated ERK1&2; Sigma #M9692), 1:250 (fixed in 8% paraformaldehyde); mouse α-Wg (4D4, Developmental Studies Hybridoma Bank),1:100; mouse α-con (C1.427, Developmental Studies Hybridoma Bank) 1:250. ABC kit (Vector Labs) was used for immunohistochemical staining. Differential interference contrast (DIC) microscopy was performed using a Zeiss Axioskop 2 microscope and Openlab (PerkinElmer) software for image capture. The following secondary antibodies were used for fluorescent staining: anti-mouse Alexa488 (Molecular Probes), 1:250; anti-mouse Alexa633 (Molecular Probes), 1:500; antirabbit Alexa488 (Molecular Probes), 1:250; anti-rabbit Alexa633 (Molecular Probes), 1:500; anti-rat Alexa633 (Molecular Probes), 1:500. Fluorescent staining was visualized by confocal microscopy using a Leica SP2 confocal microscope. In situ hybridization for detection of *mib2* and *RhoGAP15B* transcripts was as previously described (Leatherbarrow and Halfon, 2009). Color and brightness of acquired images were adjusted using Adobe Photoshop.

### Site-directed mutagenesis and transgenesis

Mutagenesis of the *mib2* enhancer was performed by overlap-extension PCR (Ho et al., 1989). Mutated sequences are shown in Fig. S2 (primer sequences available on request). Transgenic flies were generated by Genetic Services Inc. (Cambridge, MA) using phiC31-transgenesis and the *attP2* landing site.

## ACKNOWLEDGEMENTS

We thank Manfred Frasch, Eileen Furlong, Alan Michelson, Hanh Nguyen, the Bloomington *Drosophila* Stock Center (NIH P40OD018537) and the Developmental Studies Hybridoma Bank (created by the NICHD of the NIH and maintained at The University of Iowa, Department of Biology, Iowa City, IA 52242) for fly stocks and antibodies. Elizabeth Brennan, Sam Hasenauer, Qiyun Zhu, and Jack Leatherbarrow helped with experiments. Steve Gisselbrecht and Michael Buck provided critical comments on the manuscript.

## SUPPLEMENTAL FIGURES

**Supplemental Figure S1:**
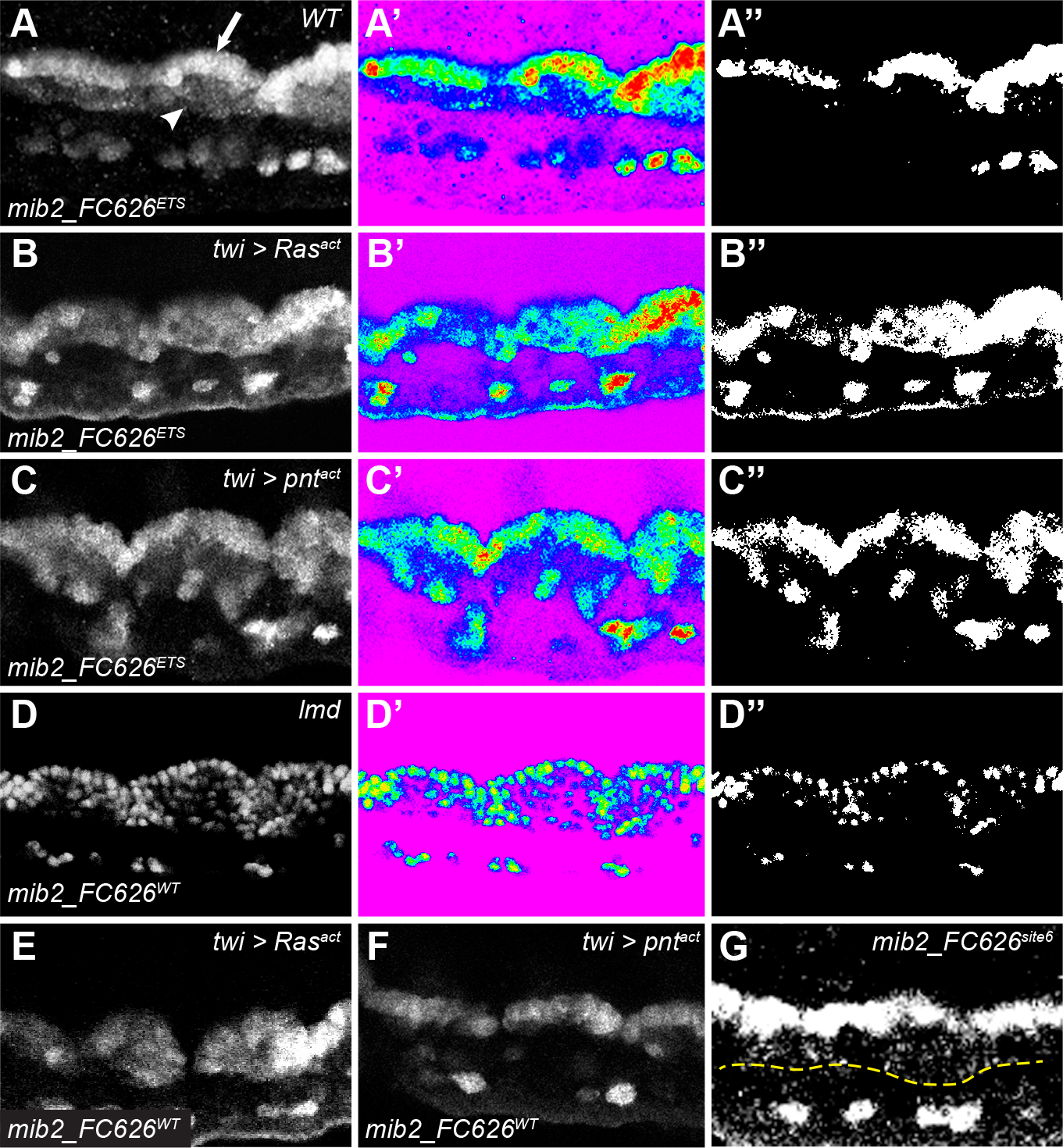
Responsiveness of the *mib2_FC626* wild type and mutated enhancers. Greyscale (A-G), pseudocolored (A’-D’), and thresholded (A’’-D’’) images of reporter gene expression in three segments of the stage 11 visceral mesoderm. Pseudocolored image show red = brighter and blue = dimmer expression. Thresholded images have had any pixels of less than one-half maximum intensity removed and the remaining pixels reset to maximum brightness. (A) *mib2_FC626^ETS^* reporter gene expression in cells specified as FCs (arrow) is stronger than the expression that expands into the FCM domain (arrowhead). (B) In an activated Ras background, however, reporter gene expression in both the native FC and FCM regions has similar strength. Reporter gene expression in the activated Pnt background (C) resembles that in the wild-type background, with FC-region expression stronger than that in the FCM region. (D) Reporter gene expression driven by the *mib2_FC626^WT^* enhancer in a *lmd* mutant embryo. Expression in the FCM domain is weaker than that in the native FC domain. (E) The *mib2_FC626^WT^* enhancer responds to pan-mesodermal expression of activated Ras and of activated Pnt (F) identically to what is observed with the longer *mib2FC_enhancer* regulatory sequence (compare with Fig. 2). (G) Weak but perceptible expansion of reporter gene expression is seen with the *mib2_FC626^site6^* enhancer. The yellow dotted line indicates the limit of the FCM domain as determined through double-labeling for the visceral mesoderm marker Biniou.

**Supplemental Figure S2:**
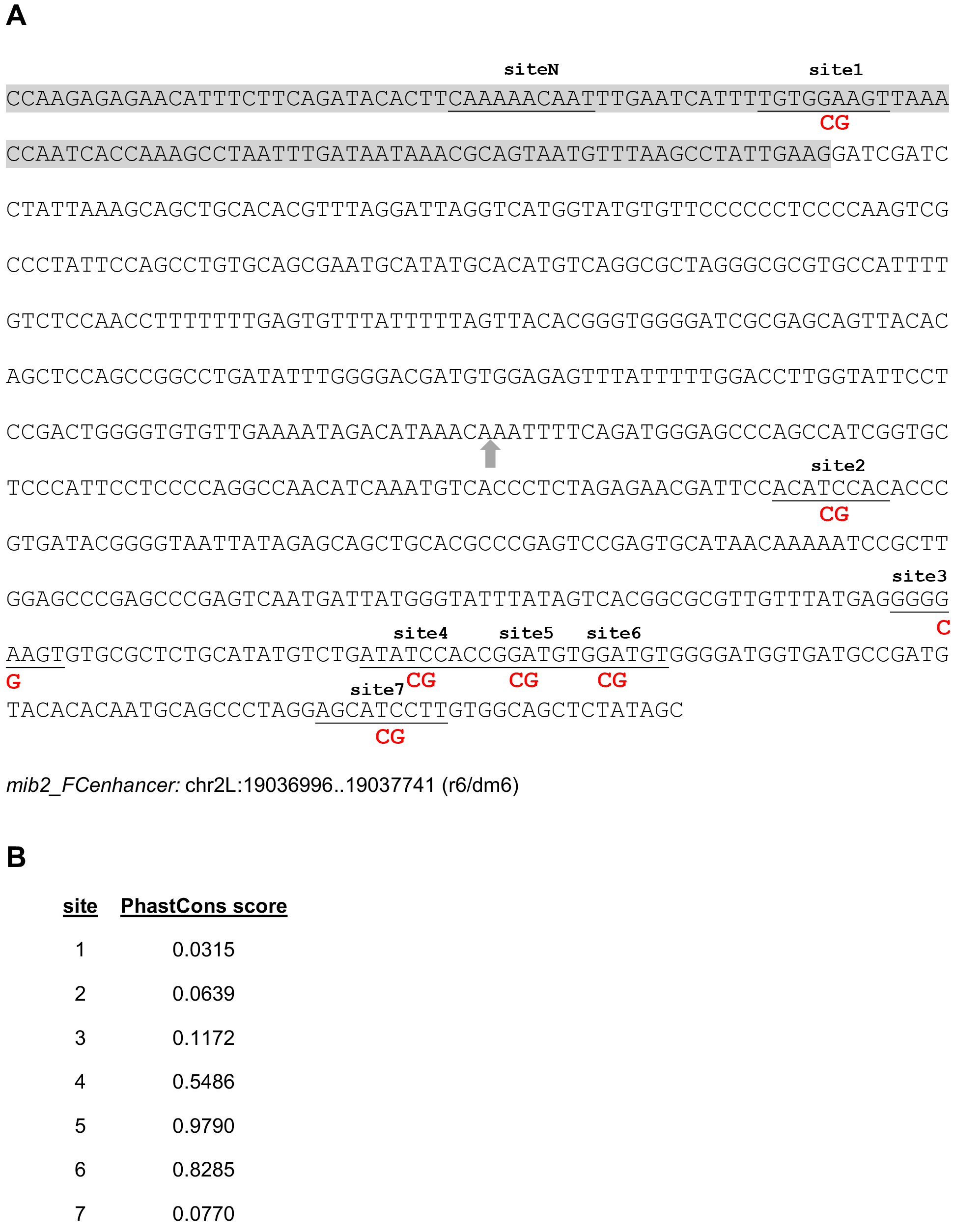
Summary of putative ETS binding sites in the *mib2_FCenhancer*. (A) Sequence of the *mib2* enhancer. ETS motifs are underlined with the substituted basepairs for the mutated enhancer shown in red. Site “N” indicates the non-canonical putative binding site derived from protein microarray data. The 5’ 120bp deletion is highlighted in gray; gray arrow marks the site of the 413 bp deletion. (B) PhastCons scores for each of the seven ETS sites (from the UCSC Genome Browser).

**Supplemental Figure S3:**
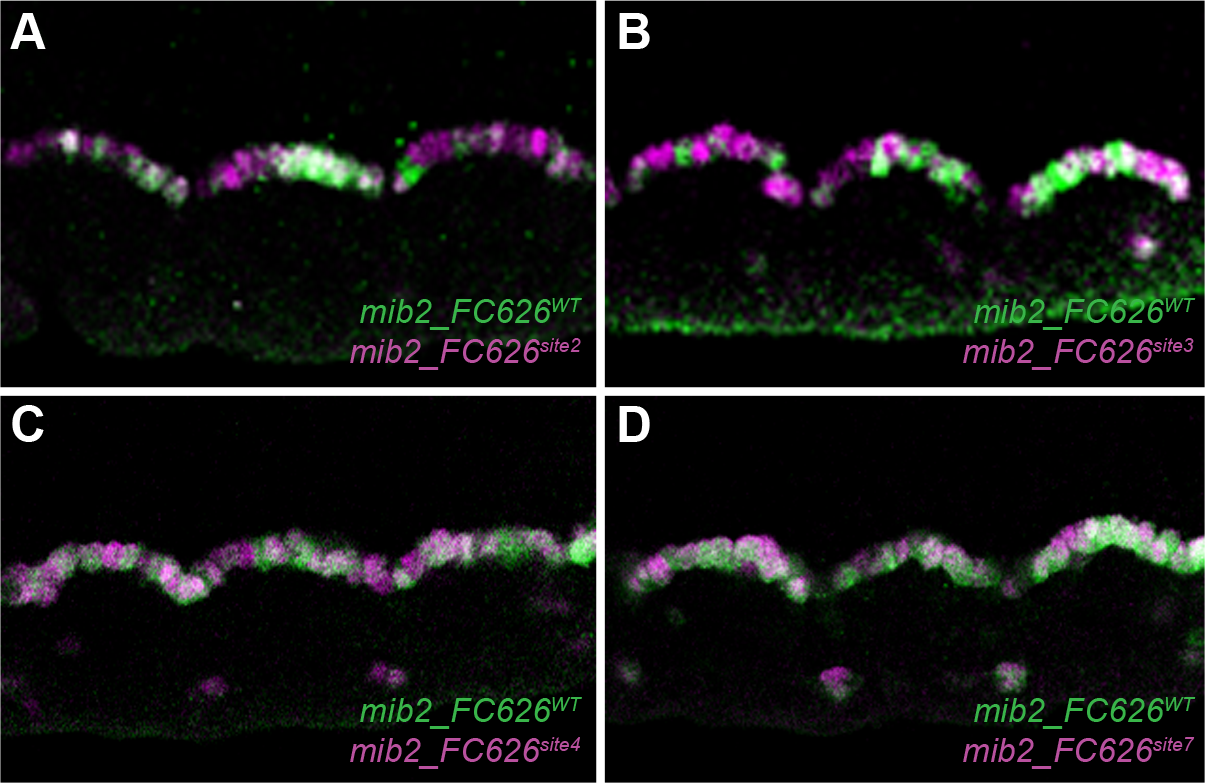
*mib2_FC626* ETS site mutations other than sites 5 and 6 have no visceral FC phenotype. Each panel depicts reporter gene expression in three segments of the visceral mesoderm, with expression driven by the wild-type mib2_FC626 in green and one of the ETS-site mutants in magenta. Mutations in sites 5 and 6 are shown in Fig. 4. Mutation of each of the individual sites site2 (A), site3 (B), site4 (C), and site7 (D) have no effect on FC expression. For effects of site7 in the nervous system, see Fig. 4L.

